# Semi-automatic stitching of filamentous structures in image stacks from serial-section electron tomography

**DOI:** 10.1101/2020.05.28.120899

**Authors:** Norbert Lindow, Florian N. Brünig, Vincent J. Dercksen, Gunar Fabig, Robert Kiewisz, Stefanie Redemann, Thomas Müller-Reichert, Steffen Prohaska, Daniel Baum

**Affiliations:** Zuse Institute Berlin, Takustraße 7, 14195 Berlin, Germany; Experimental Center, Faculty of Medicine Carl Gustav Carus, Technische Universität Dresden, 01307 Dresden, Germany; School of Medicine, University of Virginia, Charlottesville, VA 22903, USA

## Abstract

We present a software-assisted workflow for the alignment and matching of filamentous structures across a 3D stack of serial images. This is achieved by combining automatic methods, visual validation, and interactive correction. After an initial alignment, the user can continuously improve the result by interactively correcting landmarks or matches of filaments. Supported by a visual quality assessment of regions that have been already inspected, this allows a trade-off between quality and manual labor. The software tool was developed to investigate cell division by quantitative 3D analysis of microtubules (MTs) in both mitotic and meiotic spindles. For this, each spindle is cut into a series of semi-thick physical sections, of which electron tomograms are acquired. The serial tomograms are then stitched and non-rigidly aligned to allow tracing and connecting of MTs across tomogram boundaries. In practice, automatic stitching alone provides only an incomplete solution, because large physical distortions and a low signal-to-noise ratio often cause experimental difficulties. To derive 3D models of spindles despite the problems related to sample preparation and subsequent data collection, semi-automatic validation and correction is required to remove stitching mistakes. However, due to the large number of MTs in spindles (up to 30k) and their resulting dense spatial arrangement, a naive inspection of each MT is too time consuming. Furthermore, an interactive visualization of the full image stack is hampered by the size of the data (up to 100 GB). Here, we present a specialized, interactive, semi-automatic solution that considers all requirements for large-scale stitching of filamentous structures in serial-section image stacks. The key to our solution is a careful design of the visualization and interaction tools for each processing step to guarantee real-time response, and an optimized workflow that efficiently guides the user through datasets.

**Author summary:** Electron tomography of biological samples is used for a 3D reconstruction of filamentous structures, such as microtubules (MTs) in mitotic and meiotic spindles. Large-scale electron tomography can be applied to increase the reconstructed volume for the visualization of full spindles. For this, each spindle is cut into a series of semi-thick physical sections, of which electron tomograms are acquired. The serial tomograms are then stitched and non-rigidly aligned to allow tracing and connecting of MTs across tomogram boundaries. Previously, we presented fully automatic approaches for this 3D reconstruction pipeline. However, large volumes often suffer from imperfections (i.e. physical distortions) caused during sectioning and imaging, making it difficult to apply fully automatic approaches for matching and stitching of numerous tomograms. Therefore, we developed an interactive, semi-automatic solution that considers all requirements for large-scale stitching of microtubules in serial-section image stacks. We achieved this by combining automatic methods, visual validation and interactive error correction, thus allowing the user to continuously improve the result by interactively correcting landmarks or matches of filaments. We present large-scale reconstructions of spindles in which the automatic workflow failed and where different steps of manual corrections were needed. Our approach is also applicable to other biological samples showing 3D distributions of MTs in a number of different cellular contexts.

## Introduction

The segregation of chromosomes is an essential biological process and indispensable for each dividing cell. Failures in segregation can lead to defects for the emerging daughter cells as well as the entire organism. In general, missegregation is a major cause of aneuploidy and cancer. Therefore, a detailed understanding of the mechanisms of chromosome segregation is of significant biological importance.

In eukaryotic cells, chromosome segregation occurring during both mitosis and meiosis is achieved by bipolar microtubule (MT)-based spindles. The different stages of mitotic cell division are illustrated in Fig 1A. During prophase, the centrosomes start to move towards opposite sides of the cell, and a bipolar spindle mainly consisting of MTs and other associated proteins begins to form. At this point, the chromosomes begin to condense and become visible. The chromosomes then move towards the center of the cell at prometaphase. At metaphase, chromosomes are aligned at the metaphase plate and the chromatids are segregated subsequently (i.e. the sister chromatids in anaphase). Finally, the mitotic spindle is disassembled, and the nuclear membranes around the chromosomes are reformed in telophase. Quantitative analysis of MT arrangement is crucial for an understanding of spindle assembly and organization [1]. Three-dimensional reconstruction using electron tomography is an excellent tool to study the architecture of spindles with single-microtubule resolution [2]. When combined with appropriate techniques for specimen preparation, electron tomography allows an ultrastructural investigation of spindles at different stages of cell division and also a comparison of wild-type and mutant spindles, such as in the model system *Caenorhabditis elegans*.

**Fig 1.**
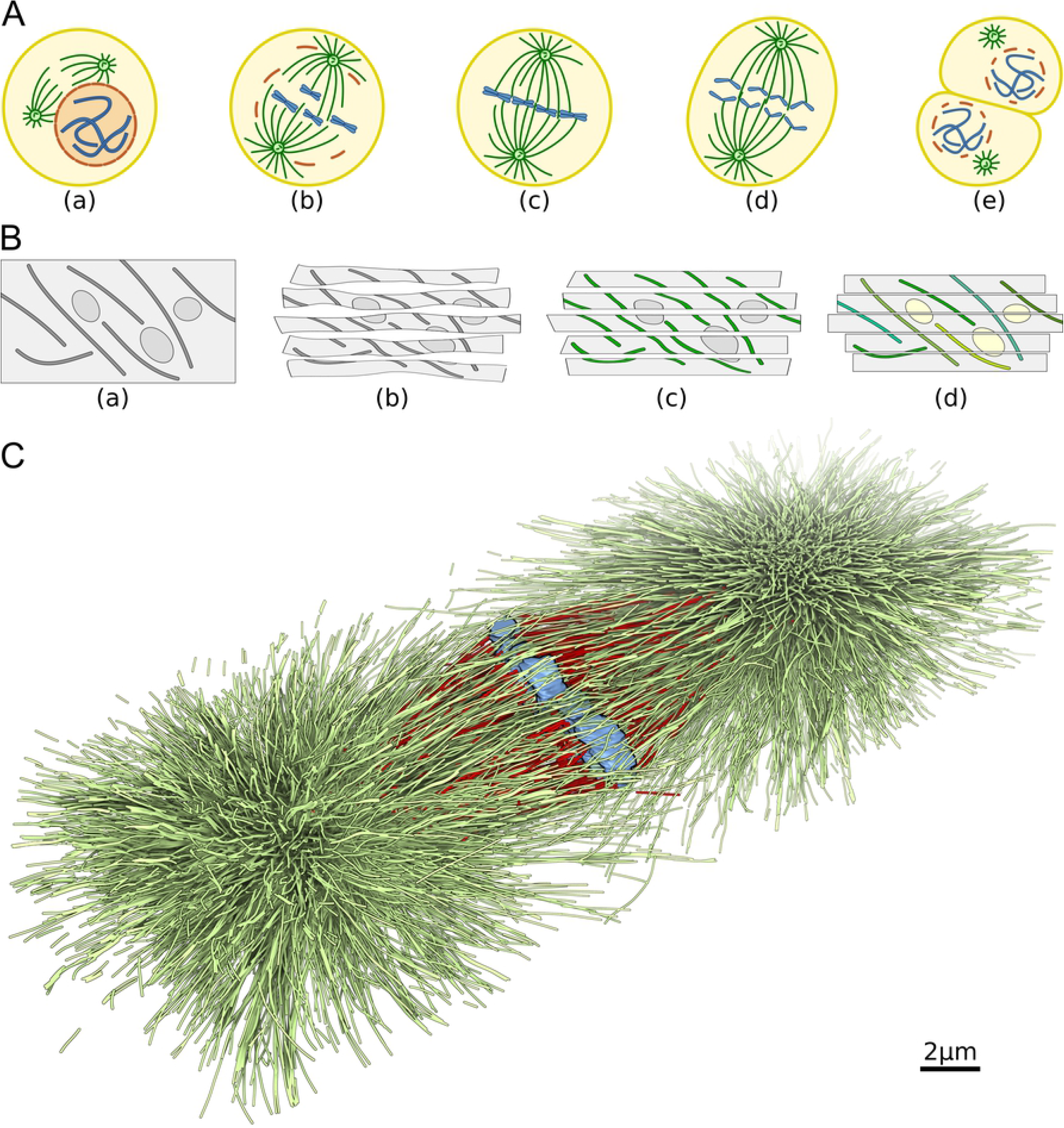
Cell division and three-dimensional spindle reconstruction. (A) Schematic illustration of mitotic stages: (a) prophase, (b) prometaphase, (c) metaphase, (d) anaphase, and (e) telophase with subsequent cytokinesis. During mitosis in animal cells, microtubules (green lines) connect the centrosomes (green circles) and the chromosomes (blue). The nuclear envelope (orange) is usually disassembled during mitosis. (B) Pipeline for the reconstruction of a mitotic spindle: (a) specimen with microtubules (lines) and other cellular components (circles), (b) physical cutting and imaging of the specimen leading to a stack of distorted serial sections (c). segmentation of cellular features within the flattened sections, and (d) stitching of tomograms to match filaments. (C) Completely aligned and stitched mitotic spindle at metaphase in the early embryo of the nematode *C. elegans* with approximately 27000 microtubules shown in green and red. The spindle was reconstructed from 23 serial tomograms (Table 1, T0265-21). Chromosomes are shown in blue, kinetochore microtubules in red [5].

In electron tomography of biological samples, imaging of serial sections is applied to reconstruct structures in 3D that are too large to image them directly. For this, samples are cryo-immobilized, embedded in plastic, and sectioned. The serial sections are imaged individually, generating one tomogram for each section. The tomograms then need to be stitched together to reconstruct the third dimension of the recorded cellular object (Fig 1B). To expand the imaged area in 2D (x/y), each serial section can consist of separate tiles that are acquired separately and then merged into a larger tomogram. Typically, such reconstructions from tiles and sections are done in ‘reverse order’, i.e. first, tiles are aligned to reconstruct a single section, called ‘montaging’, and second, the serial sections are combined to a complete stack. In this paper, we focus on the second step and assume that the first step was correctly processed. Depending on possible deformations caused by the physical sectioning and the interaction of the specimen with the electron beam during imaging, this stitching often requires a non-linear alignment. Corresponding structures in neighboring sections can be identified either based purely on image data or on segmented features.

**Table 1.** Summary of data sets as used throughout this study.

| **Data set** | **No. of sections** | **Size (GB)** | **No. of MTs** |
| --- | --- | --- | --- |
| Nocodazole-1 [25] | 17 | 8.2 | 248 |
| Nocodazole-2 [25] | 14 | 6.4 | 299 |
| T0208-1 [23] | 20 | 11.4 | 3723 |
| T0252 [5] | 24 | 13.6 | 3581 |
| T0265-21 [5] | 23 | 79.3 | 27051 |
| T0265-41 [5] | 15 | 28.7 | 11801 |
| T0265-42 [5] | 25 | 93.1 | 12205 |
| T0265 [5] | 15 | 36.4 | 8774 |
| T0275-1 [25] | 13 | 9.6 | 1511 |
| T0275-3 [25] | 7 | 9.2 | 1132 |
| T0275-5 [25] | 6 | 9.7 | 505 |
| T0275-9 [25] | 13 | 8.0 | 1289 |
| T0275-10 [25] | 14 | 8.9 | 1503 |
| T0345-11 [5] | 7 | 10.7 | 1892 |
| T0391-1 [24] | 6 | 2.8 | 246 |
| T0391-2 [24] | 11 | 10.1 | 1403 |
| T0391-3 [24] | 14 | 9.0 | 1611 |
| T0391-4 [24] | 5 | 1.9 | 671 |
| T0391-5 [24] | 8 | 3.2 | 893 |
| T0391-6 [24] | 17 | 16.9 | 2553 |
| T0391-7 [24] | 25 | 16.4 | 2212 |
| T0391-8 [24] | 11 | 9.5 | 2418 |
| T0475<sup>1</sup> | 22 | 47.6 | 4884 |
| T0479<sup>1</sup> | 29 | 79.8 | 8051 |

<sup>1</sup> Unpublished data sets (R. Kiewisz).

Typically, the spindle geometry is reconstructed in two steps (Fig 1B (c,d)). In the first step, the MTs are segmented in each serial tomogram separately using template matching. This is followed by application of an automatic tracing algorithm in combination with data verification by users [3]. In the second step, the segmented MTs are used to automatically stitch the serial tomograms to a single stack [4]. Due to the local deformations during imaging, a non-linear alignment is required that often cannot be immediately determined due to the lack of obvious image features. Thus, tomograms have to be aligned by applying a variant of the coherent point drift algorithm (CPD) to the segmented MTs. For each interface between two serial tomograms, the algorithm uses the MT ends of the two neighboring tomograms and the 3D directions at the MT ends and returns 2D landmarks as pairs of corresponding points. The MTs and tomograms are then warped using moving least squares based on the landmarks. Finally, a matching algorithm is applied on the transformed MTs to connect them over tomogram boundaries. With this approach, we previously achieved a 3D reconstruction of the first mitotic spindle in *C. elegans* embryos by electron tomography at a resolution of 5–8 nm [5], resulting in 30 tomograms with reconstructions of up to 5 GB per tomogram (Fig 1C).

Previously, the stitching was designed and implemented as a fully automatic pipeline [4], which allowed only a final verification after the complete stack was built. Furthermore, a visual verification of the complete stack was difficult. Only small manual corrections were realistic for the full stack. As an example, when the alignment completely failed for one section interface, it was impossible to create a meaningful stack. Although the following semi-automatic solution had been developed in the context of MT reconstruction from stacks of electron tomograms, it is in principle, suitable for other problems, where stacks of 3D volumes containing filamentous structures need to be aligned. The filaments are explicitly represented and used throughout the entire process to achieve high reconstruction quality with consistent filament geometry across tomograms, which seems unrealistic with generic image-based registration approaches.

A fully automatic approach to stitch such tomograms is often problematic for two main reasons. First, the data quality may be poor due to the size of the tomograms and inaccuracies during specimen preparation or microscopy image acquisition. The sections may have suffered from physical damage. As a result, they may have been deformed or ruptured, causing the alignment to fail. Second, visual validation and correction of the stitching result is difficult. An interactive inspection of each filament requires a prohibitive amount of time. With limited time, it is not obvious which subset to choose for inspection. There is also no mechanism to interactively correct the alignment and the filament matching if necessary, or to integrate user-provided information into the automatic matching process to improve it. If for some reason the automatic stitching failed between two tomograms, often the full stack could not be used anymore. Thus, in absence of options to manually improve aligned data sets of insufficient quality, the biological value of these data sets is limited.

In general, automatic methods accelerate the reconstruction of large volumes and usually operate successfully for images of expected quality. Automatic methods, however, can fail for specific cases due to unexpected data quality or underrepresented structures. In such situations, interactive tools are helpful to solve these specific cases and to allow the user to control the quality of the result. One of the most widely used semi-automatic software packages for electron tomography reconstruction is IMOD [6]. It is used in many laboratories for semi-automatic montaging and flattening of tomograms. A tool focusing on the alignment of serial tomograms of electron microscopy images was proposed by Anderson et al. [7]. The tool, named ir-tweak, provides an easy way to place landmarks with a direct real-time preview of the resulting non-linear warping using thin plate splines. We transferred this design to the alignment of serial tomograms containing filamentous structures. In contrast to thin plate splines, however, we use moving least squares for the warping. Berlanga et al. proposed a workflow for reconstructing 3D images from serial section confocal microscopy for the mouse brain [8]. Their workflow includes the flattening of the tomograms using manual landmark generation with IMOD as well as semi-automatic alignment of the section with ir-tweak. In 2012, Cardona et al. reported on a tool for investigating neural circuits from serial section microscopy images [9]. The alignment is done completely automatically. The focus of their work is the semi-automatic interaction with the image data. To explore large connectome data sets, Beyer et al. proposed a framework that enables petascale volume visualization [10]. The complete montaging and sectioning are done in real-time based on an automatically pre-computed transformation. As for Cardona et al., the tool is not designed to manually edit the transformation but to segment and explore neurites and synapses in the data. More recently, the same authors proposed an advanced tool to organize and inspect the whole segmentation process [11]. A tool for a related application, called Filament Editor, was proposed by Dercksen et al. [12]. It allows reconstructing 3D neuronal networks based on serial optical microscopy sections. The tool provides 3D visualization of the networks and image volumes as well as 2D slices and allows users to manually edit the networks. The serial tomogram alignment is computed automatically with a linear transformation. The networks that can be handled are sparse and small compared to the microtubules data sets that are the focus here.

Our goal was to stitch microtubules across a whole stack of serial-section electron tomograms in a semi-automatic way allowing for corrections during the stitching process. A number of requirements were identified for such an approach. The user should be able to influence all steps of the semi-automatic stitching process. In addition, the pipeline should be flexible such that the user can change the alignment or matching at any time. In the worst case, the user should be able to perform the whole stitching manually. In the best case, the user should be able to apply the stitching completely automatically. For cases of medium quality, minor manual interactions should support the automatic algorithms to rapidly increase the quality of the reconstruction. The whole pipeline should be supported by visualizations to verify the current state and offer options to interactively correct errors. Finally, the application should be able to interactively handle all relevant data sizes, which could be up to 30 tomograms with 5 GB each.

In this paper, we present a software-assisted workflow to address these requirements. Two related problems need to be solved: (1) the alignment problem, where the serial tomograms are transformed such that they fit together; and (2) the matching problem, where the filament structures are combinatorially connected between neighboring tomograms. In summary, we make the following key contributions:

- Design of a user-controlled workflow (Fig 2) to solve the stitching problem that integrates automatic alignment and matching. The automatic algorithms take user input into account to quickly generate and continuously improve the result.
- Implementation of a new visualization method to monitor tomogram regions that have already been visually inspected by the user.
- Development of the ‘Serial Section Aligner’, a new software tool with multi-view visualizations and interactions designed to support this workflow. A user can inspect both the images and the segmented filaments in order to validate and correct the alignment and matching.

**Fig 2.**
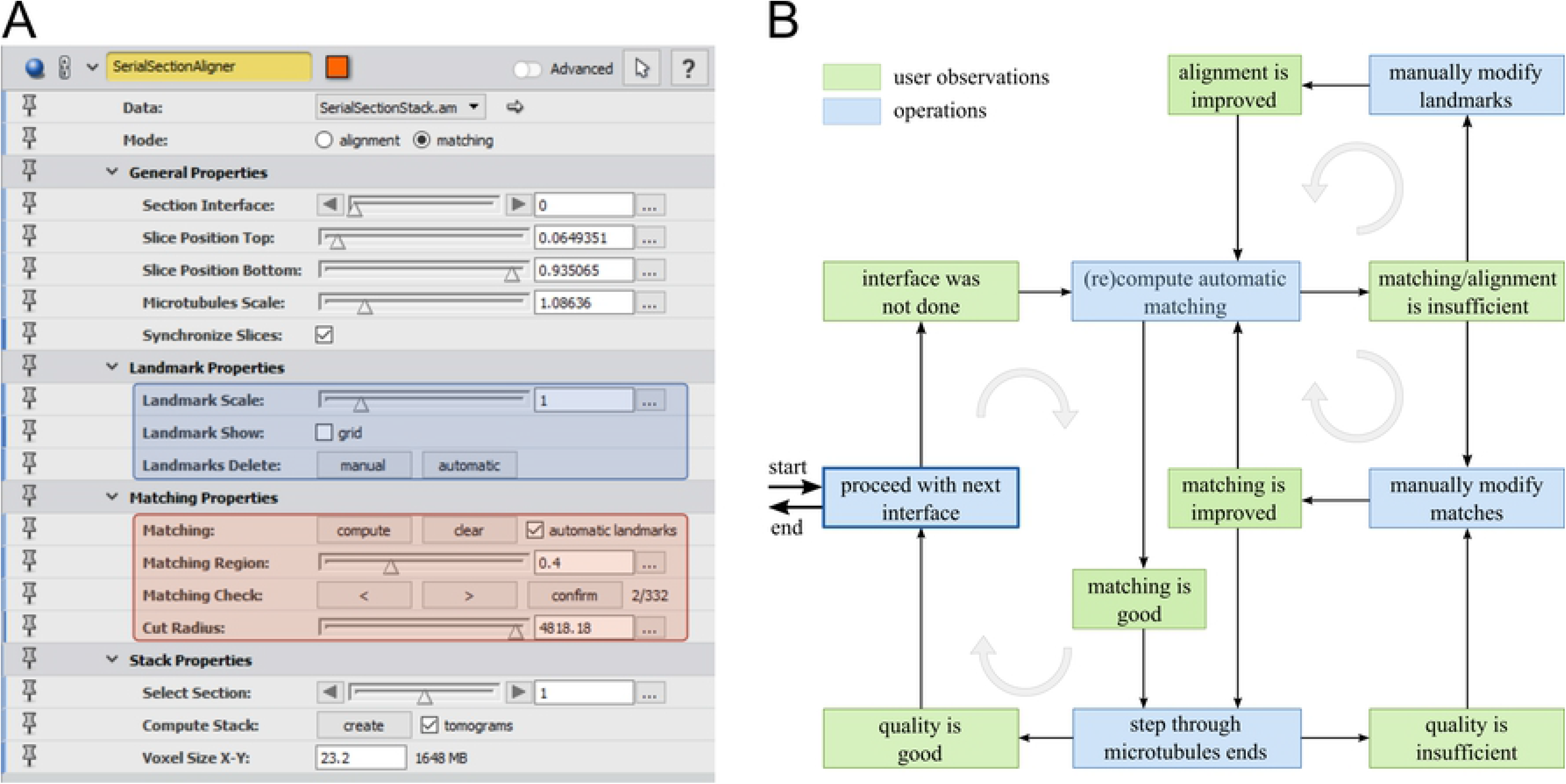
Graphical user interface and recommended workflow. (A) Graphical user interface of the aligner module. The blue part is only visible for the alignment mode, the red part only for the matching mode. (B) Recommended workflow for the stitching process. Observation tasks (green boxes) lead to operations (blue boxes). The most frequently used cycles are highlighted (gray circular arrows).

The workflow is demonstrated for serial-section electron tomography of both mitotic and meiotic spindles of high-pressure frozen and plastic-embedded material.

## Results

Based on the requirements stated in the previous section, we concluded that both alignment and matching should be first solved locally for each section interface, i.e. the cut between two neighboring tomograms. Subsequently, the full stack should be constructed from the local solutions. These conclusions lead to the following design decisions for stitching of a single section interface:

- The two tomograms of the interface should be interactively visualized with the current local alignment and matching such that errors can be quickly detected.
- Simple interaction tools should allow one to set or change the alignment with manual landmarks. Additionally, completely discarding the automatic alignment should be possible.
- Manual landmarks can be used to improve the initialization of the automatic alignment computation.
- The current matching can be changed by adding or removing matches manually. Manual matches should influence the re-computation of the automatic matching. Furthermore, the user should be able to confirm automatic matches or unmatched MT ends, which will from then on remain in the confirmed state.
- In contrast to the previous fully automatic pipeline [4], only matched filament endpoints should impact the automatic alignment in order to control the consistent microtubule geometry reconstruction across tomograms.
- The application should give immediate visual feedback about the incremental matching progress.

The stitching tool was implemented as two modules in the visualization software Amira [13]. The first module is a data class (*Serial Section Stack*) that manages the input data and the stitching state. The second module (*Serial Section Aligner*) implements the visualizations and editing operations of the stitching workflow.

### Serial section stack

A ‘Serial Section Stack’ contains the data regarding all serial tomograms of the stack including the reconstructed filaments. Assume the serial section stack contains *n* tomograms, they will be denoted by *T*_1_,…, *T*_*n*_. For the stitching of the stack (Fig 1B), we expect that each tomogram has been pre-processed, i.e distortions along the thin tomogram side, which are usually the z-direction, have been removed in a flattening pre-processing step, followed by a cropping step. Furthermore, the filaments have been segmented and verified for each tomogram. We will denote the set of filaments of tomogram *T*_*i*_ by *F*_*i*_, where each filament *f* ∈ *F*_*i*_ is a piecewise linear curve with *f* = {*p*_1_, …, *p*_*k*_} and *p*_*j*_ ∈ ℝ^3^. We further assume that the stack is always built in z-direction. We call a slice of the tomogram that lies in the x-y-plane a z-slice w.r.t. a certain z-value.

In addition to the tomograms and their corresponding filament sets, the ‘Serial Section Stack’ also stores the stitching state for each of the *n* − 1 interfaces between neighboring tomograms. A stitching state contains sets of landmarks on the tomogram boundaries and sets of matchings representing corresponding filament ends.

### Serial section aligner

The ‘Serial Section Aligner’ module provides two modes. The alignment mode allows the user to manually add, modify, and remove landmarks. The matching mode, on the other hand, enables the automatic computation of a matching in combination with manual editing and verification. In order to visualize the current state of the matching and alignment and to allow for manual interaction, the module uses a four-view visualization (Fig 3). In addition, the module has a graphical user interface (GUI) to adjust the module parameters (Fig 2A). The first part of the GUI is independent of the specific mode. It allows selecting the currently investigated section interface, adjusting visualization parameters for the interface, changing the position of the two z-slices of the neighboring tomograms, respectively, and adjusting the scale of the filament radii. The user can also choose to control quality parameters, which have a major effect on the performance. The second part of the GUI depends on the active mode, as described below. The third part allows the user to adjust and create the full stack as the final step of the stitching process. The software is available as extension package for Amira. The application is shown in the Video S1.

**Fig 3.**
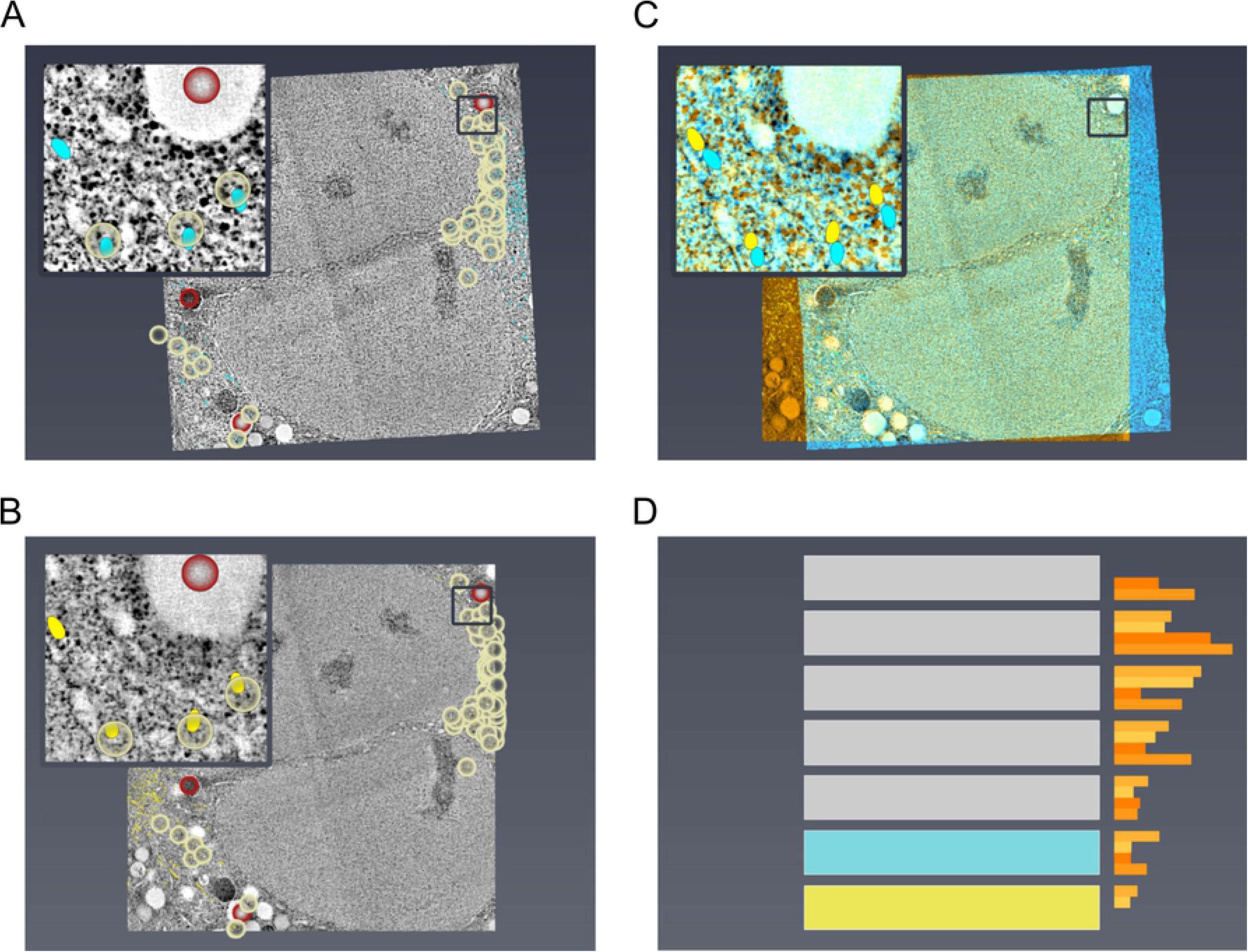
The four views of the alignment mode. (A) Z-slice of the top section with a warping preview as overlay view. (B) Bottom section. (C) Abstract visualization of the full stack. (D) Selected interface with a vertical histogram of the end point density.

#### Alignment mode

This mode is for adjusting the alignment of a pair of tomograms for a selected interface. The four views in the visualization window (Fig 3) are used in the following way. The ***top left view*** displays a z-slice of the upper tomogram of the selected interface. Accordingly, the ***bottom left view*** displays a z-slice of the lower tomogram. The user can move and zoom simultaneously in both slices to navigate. Additionally, the user can change the positions of both z-slices with two sliders (Fig 2A). The two sliders are synchronized per default; when the top z-slice is decreased by the user, the bottom one is automatically increased and the other way around. The views allow the user to identify corresponding features in the tomograms, such as vesicles, chromosomes, or membranes. To explicitly match features in the two z-slices, the user can add, move, or delete manual landmarks with mouse interactions in the two left views. Landmarks always occur as a pair of 2D points, one point for each tomogram. The points are placed such that they mark corresponding biological structures. In our visualizations, they are rendered as red circles on the z-slices. Radii of the circles can be adjusted by the user. During landmark placement, a rigid transformation is computed based on the current landmarks. The transformation globally rotates and translates the top slice, thereby ignoring distortions, such that corresponding regions are displayed for the top and bottom tomogram during navigation (panning and zooming).

In the ***top right view***, an overlay of the selected z-slices is visualized by using transparency in combination with two different colors: blue for the top slice and yellow for the bottom slice. Based on the interactions in the left views, the z-slice of the upper tomogram is warped in real time such that landmarks of corresponding locations are mapped onto each other. This allows the user to interactively evaluate the current alignment and to iteratively improve it by adding more landmarks or by modifying existing landmarks.

In addition to visualizing the tomogram data and the landmarks, the filaments that intersect the z-slices are also visualized in all three views. Filaments of the bottom tomogram are displayed in yellow and the filaments of the top tomogram in blue, corresponding to the colors of the tomogram overlay visualization in the top right view.

Finally, the ***bottom right view*** shows an abstract illustration of the whole stack and the selected interface with the pair of tomograms. The user can change the interface with a mouse click onto the illustration. Again, the same colors are used, yellow for the bottom and blue for the top tomogram. Next to the stack, a vertical histogram is shown. Each bin of the histogram shows the density of the unmatched filament ends for 25% of a tomogram extent in z-direction. At the beginning, the histogram usually shows higher densities at the interface boundaries, because the filaments are not yet matched. During the stitching process these differences decrease as more and more filament ends are matched. The plot helps the user to evaluate the overall quality but also to identify the most critical interfaces that need further investigation. A high density of unmatched ends at a tomogram boundary compared to the ends within the tomogram indicates an incomplete matching.

In addition to manual landmarks, automatic landmarks (yellow) can be generated from matched filaments. This approach is described in the following section. The user can also remove and manipulate these landmarks. However, if the automatic landmark placement is activated, these landmarks will be updated by changes in the matching.

#### Matching mode

In this mode, the user defines the matching between the filaments of the two tomograms. This is supported by four views and the possibility to automatically compute the matching in combination with manual editing (Fig 4).

**Fig 4.**
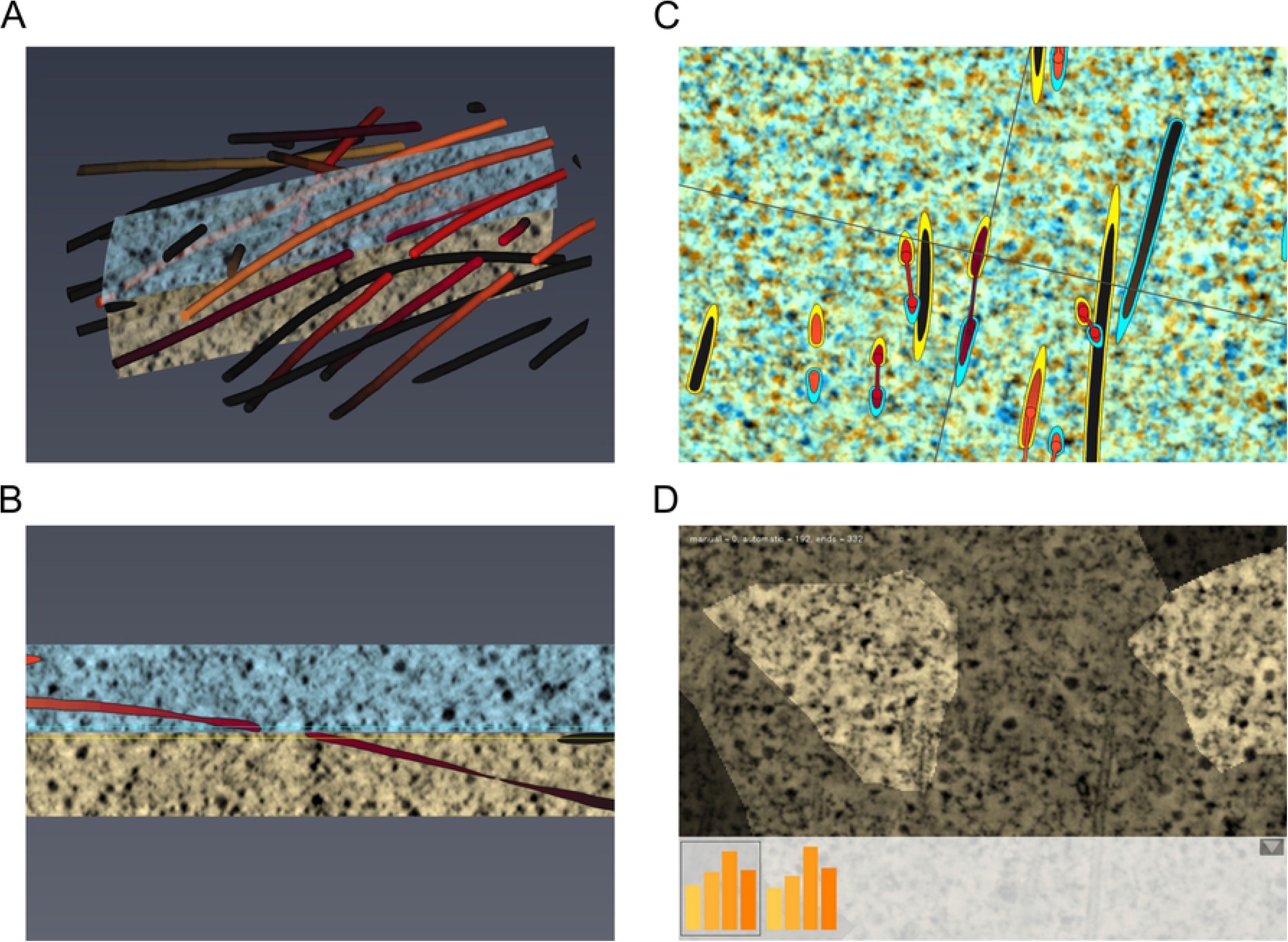
The four views of the matching mode. (A) Local 3D view. (B) Orthogonal side view. (C) Warped overlay of the bottom and the top section. (D) Matching state map of the microtubule ends of the bottom section.

In the ***top right view***, again an ***overlay*** of the z-slices is visualized together with the filaments. The filaments are colored according to the matches. Automatic matches are depicted in different shades of red and manual ones in shades of green. The colors are taken from ColorBrewer.org [14]. Unmatched filaments are visualized in black. Because a filament can be matched at both ends, the color fades out from one end to the middle of the filament to avoid ambiguous coloring. In addition to the coloring, a match between two filaments is visualized by a line, which also fades out accordingly. To keep track of which filament corresponds to which tomogram, a colored border around each filament is drawn. Again, blue is used for the filaments of the top tomogram and yellow for the ones of the bottom tomogram. The user can select two filament ends of the two tomograms with the mouse to either add a manual matching or to remove it.

To facilitate the matching decisions, the overlay view is supported by a ***side view*** on the ***bottom left*** and a ***3D view*** on the ***top left***. During navigation, the three views are simultaneously updated. To keep track of the side view, two lines are displayed in the overlay view that illustrate the position and orientation of the side view. The user can select an arbitrary filament of the bottom tomogram in the overlay view, and the side view is immediately aligned such that this filament is centered and contains the tangent of the filament. The side view enables the user to evaluate the slope of matched or unmatched filaments. It also helps detecting distortions that prohibit an automatic matching and help define manual matches for such cases. Similar to the lines in the overlay view that depict the positions of the side view, two lines in blue and yellow depict the position of the z-slices. The 3D view shows all MTs of a local region around the center of the overlay view. In this view, the user can zoom, pan, and rotate the scene. To avoid visual clutter, the region is cut by a sphere with a user selected radius. Similar to the side view, the 3D view shows a part of the tomogram within the cutting sphere as a transparent slice. The 3D view is particularly helpful to investigate the shape of multiple filaments at once.

At any time, the user can trigger an automatic (re-)computation of the matching based on the current alignment. At present, the matching is computed using the algorithm described by Weber et al. [4]. However, this part works as a black box for any suitable filament matching algorithm. The algorithm takes as input the transformed 3D positions of the filament ends with approximated 3D directions of the two neighboring tomograms. Furthermore, the algorithm receives information about all matched filament ends that should not be touched anymore. If the user checks ‘automatic landmarks’ in the GUI (Fig 2A), the ends of matched filaments will be used to generate automatic landmarks for the alignment of the tomograms. A validation stepper is embedded in the GUI and is accessible via keyboard shortcuts. It allows the user to step through all filament ends in the matching region of the bottom tomogram. The stepper automatically centers the views to the currently selected filament end. The user can confirm the current state of a filament end, whether it is matched or not. The stepper will not display confirmed ends again. All filament ends with manual matches are confirmed by default. When all filament ends of the bottom tomogram have been confirmed, the states of the ends of the top tomogram are implicitly also confirmed. For this reason, it is sufficient to observe the filament ends on only one side of the interface. Due to time restrictions, users are often not willing to confirm all filament ends, but only a fixed number such as 100–200. To prevent that these filament ends are not distributed equally over the whole tomogram, the order of the ends is shuffled at random. In addition to stepping through the filament ends, the user can also directly select a filament end of the bottom tomogram in the overlay view and confirm its state. To keep track of the state of open and confirmed ends, the GUI always displays their numbers.

The last view, the ***bottom right view***, is used to display the current state of the matching process. It shows the z-slice of the bottom tomogram, subdivided into Voronoi regions of the filament ends in the matching region. Regions that correspond to filament ends for which the user has confirmed the state are displayed in a bright color. Automatically matched ends that were not explicitly confirmed by the user are displayed less bright. The remaining regions, containing all unmatched, unconfirmed filament ends, are displayed in a dark color. The map always displays the same view for the bottom tomogram as in the overlay view. By selecting a region in this map, the stepper jumps automatically to the corresponding microtubule end. Hence, the map can help to evaluate the automatic matching and to steer the manual investigations. Especially for large data, regions in the tomogram that need further attention can thus be identified quickly.

Within the fourth view, again the end point density for the current interface is shown. The histogram consists only of four bins, two for the upper 50% of the bottom tomogram and two for the lower 50% of the top tomogram. The leftmost histogram shows the current state, while the others show the temporal evolution. A new histogram is added after each recomputation of the automatic matching. With this, the user can quickly evaluate the impact of the previous manual editing.

#### Construction of stacks

When the filament ends of all section interfaces have been matched adequately, the matchings can be used to warp all tomograms of the whole stack and create a merged volume. For this, one tomogram is selected as base tomogram. Hence, this tomogram will not be warped. The tomograms above and below the base tomogram are warped based on the landmarks of the corresponding interfaces. The warped tomograms are then resampled into a single volume containing all tomograms of the whole stack. The warping is done similarly for the filaments. Matched filaments are joined over the section interfaces. Each point of a filament receives an integer attribute that stores the original tomogram ID of the point. If the quality of the tomograms and the distribution of the filaments is more or less uniform over the full stack, the middle tomogram should be ideal as a base section to avoid accumulating the warping error over more interfaces than necessary. Otherwise, one should select a tomogram close to the middle of the stack with the best quality and filament distribution.

The GUI has a dedicated parameter section for the creation of the full stack. A checkbox allows the user to create only the stack of the filaments without the aligned tomograms, which is much faster. Choosing a lower resolution for the x- and y-directions of the tomograms is another way to increase speed and reduce memory consumption. Both options help to quickly generate a preview of the stack or create results on systems that do not have enough RAM to merge the tomograms at full resolution.

### Recommended workflow

For applying the ‘Serial Section Aligner’, we recommend the following overall workflow (Fig 2B). After creating the ‘Serial Section Stack’ object and detecting the correct tomogram order, the section interfaces will be processed one after another. For each interface, the sections should be sufficiently aligned and the filaments matched such that it should not be necessary to revisit the interface. This is recommended, because frequent switches between section interfaces can be time consuming due to repeated loading of tomograms. When the quality of the tomograms is good and the filaments are distributed well over the whole tomograms, the user should directly switch to matching mode and check the automatic computation of the matching. If both matching and alignment look good, the user should step over a fixed number of filament ends and confirm or possibly correct them. Finally, if in some regions there are very few or no filaments, the user should consider adding a few manual landmarks in those regions.

If the automatic matching failed completely, the user should clear the matching. Now, there are two possibilities to continue, which can also be combined. First, the user sets a few manual landmarks in alignment mode and reruns the automatic matching. Second, the user manually matches a few filaments, which also results in new landmarks, and reruns the automatic matching. Usually, the second option is only reasonable if the sections are almost aligned. As soon as the automatic matching seems to produce reasonable results, an iterative process starts with correcting or adding manual matches in combination with an automatic re-computation. Usually, small manual corrections help to improve the whole region in the neighborhood with the automatic matching. It is recommended to look for the same unique patterns of filament ends in both sections by moving the z-slices. An example is shown in Fig 5A,C, where the automatic matching failed locally, and the placement of two manual matches helped to improve the automatic matching for the complete region (Fig 5B,D). Finally, the same procedure is followed as described above for a direct successful automatic matching.

**Fig 5.**
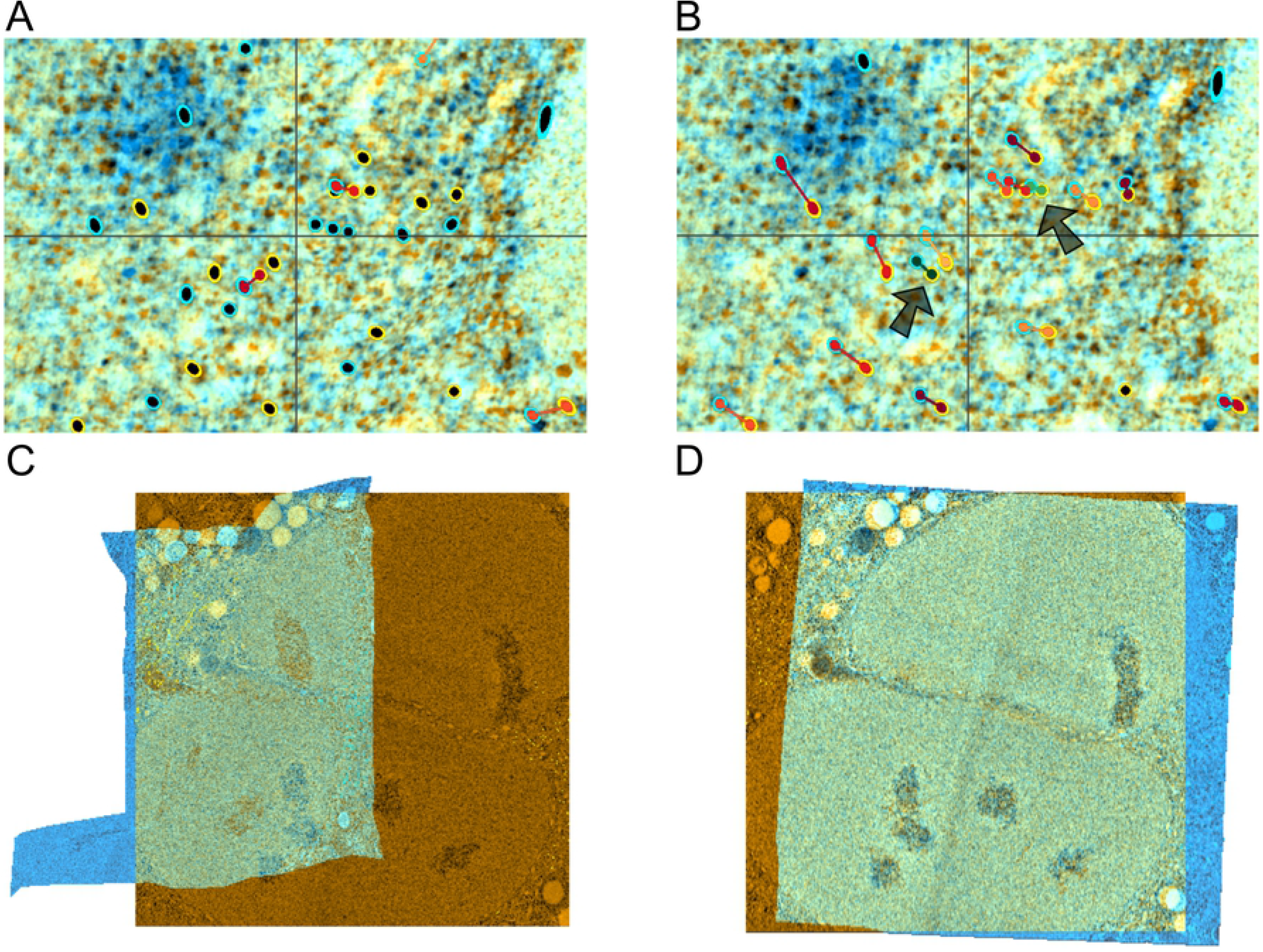
Fully automatic approach *vs.* semi-automatic workflow. (A) Failure of the automatic matching in the given region, but one can clearly see the matching patterns for the MT ends. According to this, a shift of the top tomogram (blue) up and slightly to the right would be required (Table 1, T0265-42). (B) By adding two manual matches (in green) the automatic matching for the complete region worked. (C) Alignment result of the original automatic pipeline in a *C. elegans* prometaphase data set of one section interface (Table 1, T0345-11). (D) The problem was solved by adding a few manual filaments matches.

When the direction of the filaments is highly distorted, it might be useful to either completely deactivate the automatic landmark placement by the checkbox in the GUI or forbid the landmark computation for a certain filament match. The latter can be done with a keyboard shortcut for the selected matched filament end. Already placed automatic landmarks can be removed in the alignment mode. In case the automatic landmark placing is disabled, mainly modifications of manual landmarks can improve the automatic matching.

### Technical aspects

#### Stack data structure

As described above (see section *Serial Section Stack*), the serial section data is handled in a stack data structure. If *n* is the number of sections, then the stack stores *n* tomograms *T*_*i*_ with their corresponding filament data sets *F*_*i*_. The data sets are stored in the file system. The stack data structure loads tomograms and filaments on demand and hands them over to the ‘Serial Section Aligner’ module. During alignment and matching, only two tomograms are loaded into RAM at the same time.

For each of the *n* − 1 section interfaces between consecutive tomograms, a set of landmarks and a matching is stored. As briefly described above, landmarks always occur as pairs of 2D points in x-y-space, one point for each of the neighboring tomograms. The points mark corresponding locations on the two sides of an interface and define an x-y-transformation that warps one tomogram into the space of the neighboring one such that the tomograms and filaments are aligned. Because the tomograms are thin with a flat boundary in z-direction, after the pre-processing step, warping is only defined as an x-y-transformation, ignoring z. In addition to the pair of points, each landmark has a type attribute that stores the source of the landmark. The source can be manual or automatic. As described earlier, these types will be colored in red and light yellow, respectively.

A matching is stored as a set of quadruples. A match between *F*_*i*_ and *F*_*i* + 1_ is stored as (*a*,1,*b*,0), where the end of filament *a* ∈ *F*_*i*_ is matched with the start of filament *b* ∈ *F*_*i* + 1_. Similar to the landmarks, each match has a type attribute that is either manual or automatic.

Filament end states are stored for the bottom tomogram of each interface to maintain the matching state. A filament end state can be either confirmed or undefined.

#### Slice rendering

The rendering of tomogram slices is essential for the visual assessment of the filament alignment and matching. Our tool supports fully interactive visualization of top- and side-view slices for both original and warped tomograms.

Let us first consider the case without warping. The cost for rendering a slice that is computed in tomogram space scales with the tomogram size. If the dimensions of the tomograms are large, it can be difficult to achieve interactive frame rates. The screen, however, has a limited number of pixels. In most cases, either only a part of the slice is displayed, or it is far away from the camera so that the size of a voxel is much smaller than a pixel. Hence, small details will not be visible, and it would be inefficient to render a slice at full resolution in tomogram space. Instead, we render slices in view space. For this, in the first step, we compute for each pixel of the view the 3D position within the slice. This is done by computing the view ray through the pixel and detecting the intersection of the ray with the slice plane. Additionally, it is checked if the intersection point is located outside the bounding box of the tomogram. In the next step, we evaluate for each point inside the tomogram the intensity value using trilinear interpolation. Both steps are performed on the CPU using OpenMP, because tomograms may not fit into GPU RAM. Finally, a screen space-filling quad is rendered that is textured with the computed intensity values. With this approach, the number of memory accesses into the tomogram is limited by the number of pixels of the screen. Although we observed that rendering is interactive even for large tomograms and many cache misses, the user can explicitly switch between different slice qualities. For high quality, the full view resolution is considered, for medium quality, half the resolution, and for low quality, only one fourth of the view resolution. This can further increase the interactivity, especially on large screens.

Next, let us consider the warped case for the overlay views of the tomograms. Again, the 3D position of the slice for each pixel is computed in the space of the bottom tomogram, but without the check whether a point is located inside or outside the tomogram. To get the correct intensity values of the top tomogram in the space of the bottom tomogram, the positions need to be transformed into the space of the top tomogram. This is done usingthe moving least squares approach by Schaefer et al. [15]. For a large number of landmarks, real-time rendering of the slices might not be possible, especially for changing landmarks. However, typically the number of landmarks is much lower than the number of voxels within the slice. Furthermore, local warping results cannot be observed if the slice is far away from the user. For this reason, we apply a 2D grid in screen space. Then, for each grid point, the corresponding 3D point on the slice is detected and only the grid points are warped. For all points within a grid cell, bilinear interpolation of the warped cell is applied to compute the warped position (Fig 6). This accelerates the rendering of the warped slice, so that it can be interactively updated during landmark or slice changes. Additionally, it creates a kind of level of detail: the correctness of the warping increases the closer the user zooms into the slice. Since the warping affects only the x-y-space of the tomogram, it is sufficient to use a 1D grid for side views.

**Fig 6.**
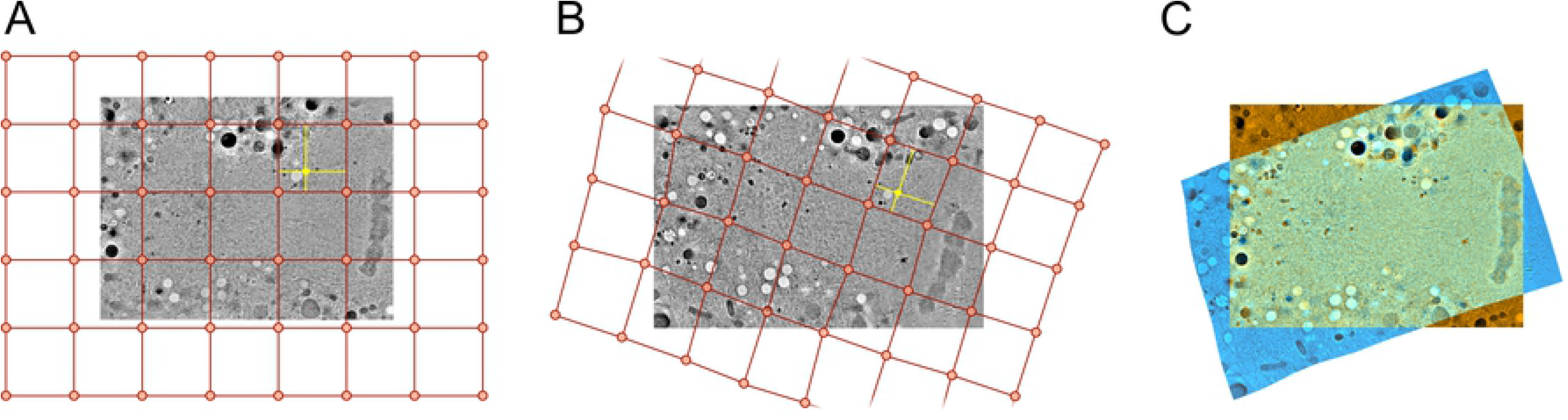
Warping of the z-slice of the top section. (A) Generation of a 2D grid in the space of the bottom section that fills the complete view. (B) Transformation of the grid to the space of the top section. (C) Approximation of all visible points of the bottom section by a bilinear interpolation in the transformed grid. The intensity value is then captured and blended with the bottom section.

We can further increase the rendering performance for the warped slices using an approximation of the moving least-squares approach. In its original implementation, all landmarks are used to transform a point. However, since the influence of a landmark decreases with increasing distance, it is sufficient to detect the closest k landmarks to transform a point. To do this in an efficient way, all landmarks are stored in a grid data structure.

#### Landmark rendering

Each landmark is visualized in the alignment mode as a circle on the z-slice (Fig 3A,B). The circles have a constant size in screen space with a scaling parameter. This seems to be superior to object-space radii, because the visible circle size remains constant across zoom levels, which simplifies user interaction. For each circle, a single vertex is generated that is then extended to a quad in the geometry shader on the GPU. The center of the quad is the center of the landmark, and the sides equal the diameter of the circle. Then, in the fragment shader, the corresponding position on the quad is inserted into the implicit circle equation to check whether the fragment is inside or outside the circle. All fragments inside the circle are colored according to the landmark type. In addition, the transparency of the color is increased for fragments close to the center of the circle. This allows the user to see the image feature that is related to the landmark.

#### Filament rendering

As mentioned in Section ‘Serial Section Stack’, each filament is represented as a piecewise linear curve, given by a set of points *p*_1_,…, *p*_*n*_. Each filament in our reconstructions is represented by a sphere at each point p_i_ and a cylinder between two points *p*_*i*_ and *p*_*i* + 1_ with constant radius of both the spheres and the cylinders. To display the filaments within the selected slices in the tomogram, only the intersection of the spheres and cylinders with the slice is visualized. To avoid an expensive triangulation of the spheres and cylinders with additional intersection computations, the rendering is based on the ray casting of quadrics proposed by Sigg et al. [16]. Each sphere is represented initially as a single 3D vertex. A geometry shader is used to create a quad from this vertex that encloses the sphere after projection. Ray casting is performed in the fragment shader. Similarly, for each cylinder, two 3D vertices are created for the start and end points, respectively. Then in the geometry shader, a quad is created that encloses the cylinder after projection. Finally, ray casting is used in the fragment shader to find the intersection of the view ray with the cylinder.

To quickly update the filament visualization of the warped tomogram in the overlay view, all spheres and cylinders that intersect the slice are detected in a first step. Then, only these spheres and cylinders are warped and visualized in a second step.

Both, filaments and landmarks, are rendered first in a frame buffer. During the rendering on the slice, a shader is used to apply anti-aliasing to the boundaries. To do so, for each fragment at the boundary, the tangent line is computed based on an analysis of the neighboring fragments in the four main directions. The line is used to compute a blending value for the fragment. Furthermore, for close-up views on the filaments or landmarks, the line is used to create a pixel-wide anti-aliased black silhouette. This procedure drastically increases the visual quality and helps to distinguish between slice data and filaments or landmarks.

#### Filament end state map

In matching mode, a map depicts which filament ends are already confirmed, which ends are automatically matched but not confirmed, and which ends are unmatched and not confirmed (Fig 7). This map is generated and updated in the following way.

**Fig 7.**
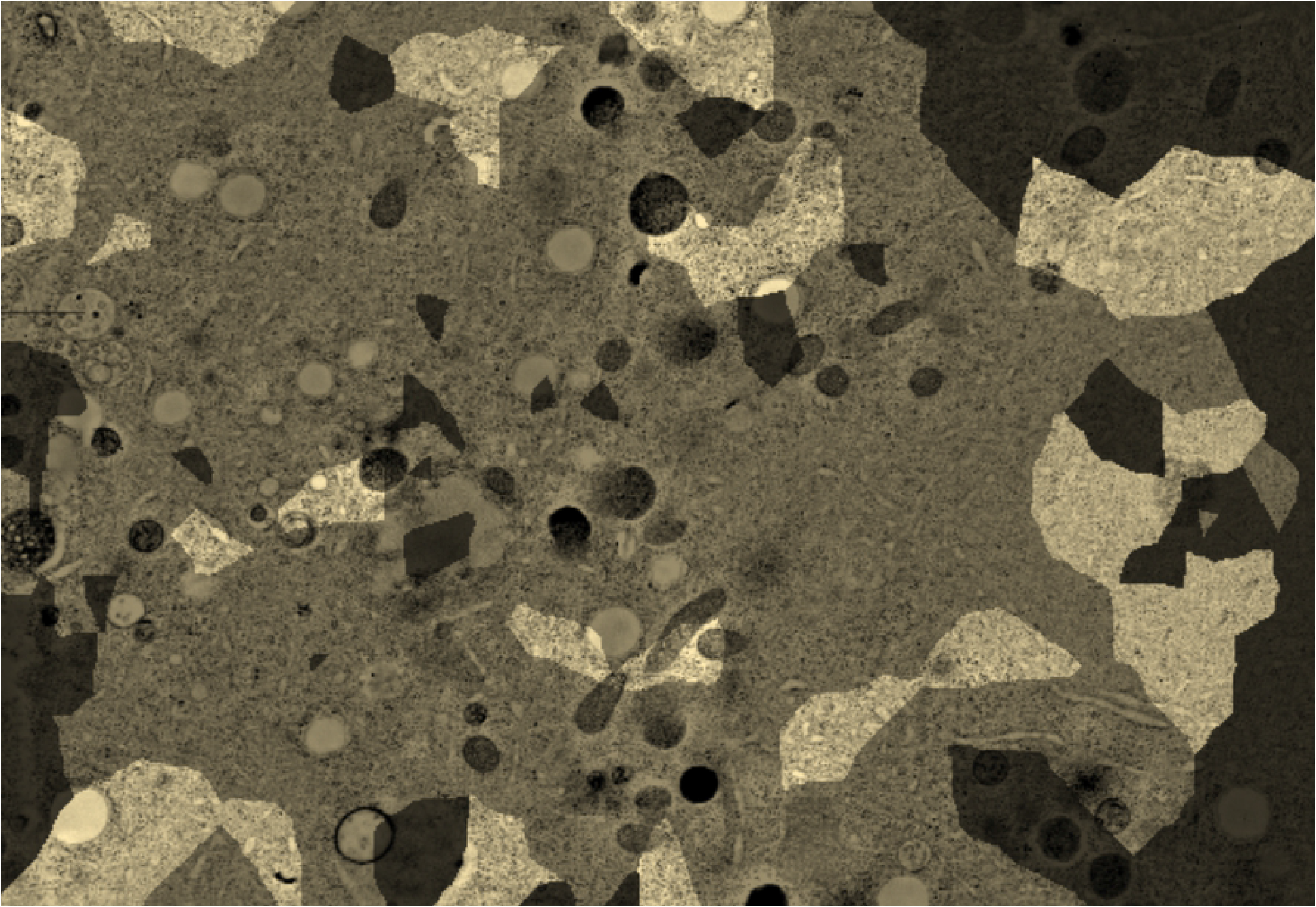
Filament end state map. Dark regions correspond to unmatched unconfirmed filament ends, bright regions to confirmed ends, and the ends with medium brightness are automatically matched but not confirmed.

First, a discrete 2D scalar field is created that has the same boundary in the x- and y-directions as the bottom tomogram. Then for each sample position of the scalar field, the closest distance to all filament ends in the matching region of the bottom section is computed. The identifier of the closest filament end is stored at this position in the scalar field. Hence, the scalar field is a discrete representation of the Voronoi diagram of the filament ends in the matching region. Note that this map needs to be updated only when the user changes the matching region.

Then the z-slice of the bottom section is rendered as described above. However, during the detection of the intensity value in the tomogram for a pixel, additionally the closest filament end is determined from the pre-computed Voronoi diagram. Then the state of the end is used to modulate the brightness of the intensity value, as described previously in the ‘Matching Mode’ section. An illustration of the filament end state map is displayed in Fig 7.

### Matching and alignment examples

Using the described tools, more than 20 data sets were assembled so far. The data sets were obtained from different specimens, different types of spindles (i.e. mitotic *versus* meiotic ones), and different stages of cell division, as well as from both wild-type and mutant samples. In Fig 8, four of these spindles are shown as reconstructed tomogram stacks with stitched MTs. These examples demonstrate the structural diversity of microtubule-based spindles.

**Fig 8.**
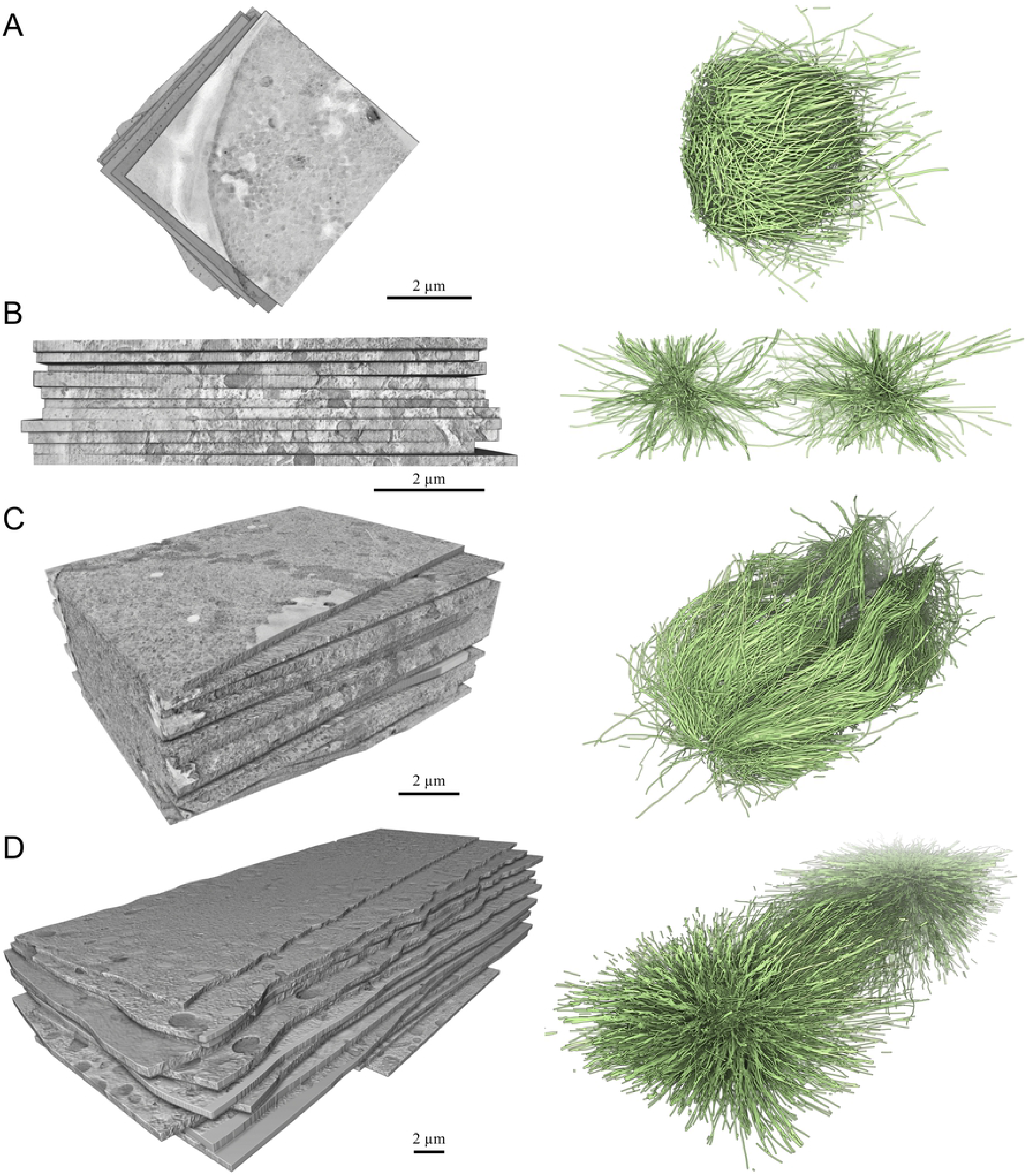
Three-dimensional reconstructions of spindles from different biological samples. Tomographic stacks are shown on the left, 3D reconstructions of the corresponding spindles on the right. (A) Anaphase I of oocyte meiosis in *C. elegans* (Table 1, T0208-1) [23]. (B) Anaphase I of spermatocyte meiosis in *C. elegans* males (Table 1, T0391-2) [24]. (C) Metaphase in mitotic HeLa cell (Table 1, T0475). (D) Metaphase of the first mitotic division in the *C. elegans* embryo (Table 1, T0265-21) [5].

In the following paragraphs, three challenging datasets are described in detail. For the first two, experimental problems had to be tackled, such as damaged sections or poor image quality. The third example represents the largest spindle that has been reconstructed so far. We used it as benchmark for our system.

#### Mitotic spindle at prometaphase

In this phase of mitotic cell division in *C. elegans*, numerous MTs emenate from the centrosomes and enter the pronuclei through the fragmented nuclear membranes to make contact to the chromosomes. The sample stack contains a region around the pronuclei and consists of 7 sections with 300–500 MTs per section (Table 1; T0345-11; [5]). Due to the field of view with the nucleus in the center, most of the MTs are close to the image boundaries in x-y. Hence, the original, fully automatic workflow failed (Fig 5A,C). None of the section interfaces could be aligned. The reason was most likely the uneven distribution of the MTs. With a few manual landmarks, often only 3–5, the automatic matching worked quite well. If the global transformation between two sections was moderate, the same result could often be achieved with a few manual matches. Finally, for each section interface, further manual matches or corrections were applied to achieve the desired quality. An experienced user processed a section interface in approximately 10 minutes. The complete stack could thus be constructed within one hour. About 25–35 % of the matches were placed manually. Substantial manual effort was especially necessary close to the section boundaries in x and y.

#### Mitotic spindle at anaphase

This stack consists of a half spindle in early anaphase in *C. elegans* (Table 1; T0265-42; [5]). The chromatids split and move towards the poles. The data set includes a half spindle with centrosome. It consists of 25 sections with 1100–4200 MTs per section. However, one section in the middle of the stack got damaged during sample preparation, and the spindle half of the spindle could, therefore, not be recorded for this section. The automatic alignment of the damaged section failed, which also affected the following sections, such that the stack got distorted in z-direction. With our tool, it was possible to manually steer the alignment and create a reasonable matching for the uncorrupted data. The stack could be completely constructed. Up to 20 manual landmarks and approximately 100 manual matches were used to compute additional 1100 automatic matched MT ends. The MT distribution for the 3 sections around the damaged section is 4176, 2134, and 3774, where the one in the middle is the damaged section. It was possible to handle the two interfaces above and below the damaged section within one hour.

#### Mitotic spindle at metaphase

This stack consists of a *C. elegans* spindle in metaphase (Table 1; T0265-21; [5]). The full spindle shows the chromosomes located in the center of the cell and attached to a subset of MTs, called kinetochore MTs. Similar to the anaphase spindle, the data set includes the complete spindle with centrosomes. The spindle consists of 23 sections with 900–3900 MTs per section. In contrast to the previous two examples, the quality of the sections was good. The automatic algorithms worked quite well. We chose this example mainly to test the performance but also to check if the automatic workflow within the new tool works for the complete spindle.

### Performance

#### Memory usage

During alignment and matching, the tool requires memory for the two tomograms of the selected interface. The alignment and matching can be performed on an advanced desktop system. For our largest data sets, approximately 10 GB were sufficient. For the creation of the stack in full resolution, approximately (*n* + 1) ∙ *k* memory is required, where *n* is the number of sections and *k* the size of the largest tomogram. For our cases, 150 GB was sufficient for the reconstruction of the largest stacks in full resolution.

#### Rendering performance

The interactivity of the application mainly depends on the performance of the rendering algorithms and the required internal data structure updates. The most expensive part for the visualization is the matching mode, where three views display both the bottom section and the warped top section with microtubules. Furthermore, a map depicts the end states of the MTs of the bottom section. The rendering performance depends mainly on the resolution of the views. For our tests, we used a desktop computer with an Intel Xeon i7 X5650 processor (6 cores) and an Nvidia GeForce GTX 1080 Ti graphics card. Amira was run on a screen with 1920 × 1200 pixels. The view size of all four equally-sized views together was 1400 × 1000. For small and mid-sized sections, we measured up to 25 frames per second, while for the largest data sets the performance decreased to 12 frames per second. Thus, with default desktop systems, it was possible to align and match serial sections for all our use cases with interactive frame rates. Since the most computationally expensive parts are done on the CPU, like the computation of the intensity values of the slices as well as the warping of the slices and the microtubules, the CPU represents the bottleneck for the full rendering pipeline. Compared to this, the rendering of the slices and the ray casting of the MTs and landmarks is negligible.

## Discussion

We have demonstrated that the proposed tool solves the matching and alignment of serial-section tomogram stacks for data sets of different size, number, and distribution of MTs, including the 3D reconstruction of the full stack as one single volume data set. An important design decision was that the construction of complete stacks is decomposed into smaller tasks that solve the alignment and stitching problem separately for each interface between two neighboring sections. This seems reasonable based on the assumption that the alignment and matching for one interface is independent from other interfaces. Such a workflow has advantages for both, the visual design as well as the performance.

### Visual design

Two z-slices of neighboring sections together with a blended interactive overlay preview to align neighboring sections has been successfully used for serial sections images before [7]. Our observations confirm that the approach works well and allows a user to quickly assess the alignment visually.

Due to the low signal-to-noise ratio of electron tomography data, rendering of slices seems superior to volume rendering or other 3D rendering techniques for image data. Slice rendering directly displays the intensity values, is straightforward to implement, and scales reasonably well with image data size. We doubt that 3D visualizations of the image data would help to improve the alignment and matching. Moreover, 3D techniques often require more parameter adjustments than slice rendering, such as transfer functions, and may therefore be more time-consuming to control in practice.

A stack can contain thousands of densely arranged MTs. Visualizing the MTs as 3D lines for the full stack is inadequate for inspecting the matching at section interfaces. Due to occlusion, navigation in dense sets of 3D filamentous structures is inherently difficult. However, if the 3D visualization is restricted to a certain location, it can help to match the filaments in difficult setups, because the slope of the filaments can be seen in all directions at once, which is difficult in a single slice. On the other hand, showing MT ends as overlay of two z-slices allows the user to easily detect patterns in both sections, which can be more difficult in a 3D visualization that allows arbitrary viewing angles. For this reason, we decided to provide both, a full 2D view of the filaments within the slice and a local 3D view.

For the rendering of MTs, cylinders seem to be superior to thin lines. The intersection of cylinders with slices creates ellipsoids instead of points. The shape of the ellipsoids allows one to perceive the slope of the MTs, which helps to detect corresponding ends.

The map that depicts the MT end states was inspired by real-time strategy games. Most of these games offer a map that is initially black to indicate undiscovered regions. During the game, the player discovers regions with units. The visited terrain becomes visible on the map and the black is replaced by map details. Games usually distinguish further between regions that are currently under control by the player and regions that were visited before but are currently not under control. While the latter are usually visualized a bit darker, the others show additional information such as enemies in the regions. In our case, the bottom section is initialized as an undiscovered map. Automatic unconfirmed MT ends correspond to ‘visited’ regions. The confirmed MT ends correspond to regions ‘under control’.

### Technical aspects

The design decision to restrict the workflow to processing interfaces between two neighboring sections one after the other has several advantages. Handling data and rendering of the full stack at once can be avoided. Large-scale image data visualization could be quite complex and often requires advanced out-of-core rendering techniques [10]. Since we restricted the design to a single interface, simple and efficient slice visualizations can be rendered interactively without complex data handling. The most challenging part is the real-time warping, which we accelerated by grid data structures to achieve interactive frame rates (Section Result – Technical Aspects – Slice Rendering). We expect that due to this design even larger tomogram sizes can be handled without major modifications. Because most visualizations are based on screen-space, the aligner tool should scale reasonably well with data size. The integration of alternative automatic alignment and matching algorithms is straightforward, since the design does not use any advanced data structures.

The top left view in alignment mode (Fig 3A) is rotated and translated to approximately match the warping applied to the overlay viewer. We tested two approaches for this adjustment before we decided to use the global approach. The transformation could be calculated either by the whole slice (global) or by the current view of the bottom section (local). Although the local option more closely approximates the current local transformation, it is also visually less stable and less intuitive to use, because the viewer may adjust rapidly during zooming. A global rigid transformation is more stable and was sufficient in all test cases to display the corresponding regions in both views.

### Practical aspects

Especially the interactive visual feedback that shows the current state of alignment and matching for the section interfaces was important to assess the quality of the results. For the alignment, we considered our initial design as sufficient. For the matching, however, we concluded that a stepping tool to confirm MT ends and the visualization of the MT end state map would be needed.

For large number of MTs, it is impossible to check all MT ends per section interface. To achieve a good distribution over the whole section interface, we decided to compute a random order of the MT ends for the stepping tool. Another idea was to select an order where the next MT end is always farthest away from all previously confirmed MT ends with periodic boundary conditions. This could potentially generate a distribution that would more quickly cover the entire section. It would, however, require more computation time and seemed irrelevant for our use cases. Although the stepping tool alone already helped validating the matching, we found it difficult to estimate the overall matching quality for the entire interface. For this reason, we introduced the map that depicts the state of the MT ends to give an impression of the overall progress.

Since it is unfeasible to check all MT ends for large numbers of MTs, it is expected that some errors will remain in the final matching and hence the final alignment. Therefore, a trade-off has to be made between efficiency and accuracy. Checking a certain number of MT ends (about 10–25) after being visually satisfied with the alignment should give the user an idea about the expected quality of the result According to previously published data, approximately 5% of MTs needed to be corrected [5]. These corrections included the manual tracing of undetected MTs, the connection of MTs and deletions of tracing artifacts (for example, membranes of vesicles).

Application of our approach indicated that the position and orientation of the side views are very helpful. The lines primarily help in two ways. First, they provide a visual link between the overlay view and the side view, and second, the selected MT end is highlighted by aatcross (i.e. where the two lines intersect) and thus easier to inspect.

Originally, the automatically computed landmarks were maintained separately from the matching process. The coherent point drift algorithm could be applied completely independently from the matching, as in the fully automatic pipeline [4]. We observed that this separation was confusing in practice. The landmarks are computed based on the geometry of the MT ends in the matching region and thus conceptually belong to the MT ends. We observed that users either accepted these landmarks or rejected them, because the alignment completely failed. They rarely modified the automatic landmarks. For this reason, we decided to move the original alignment into a hidden pre-processing step before the automatic matching, which is only computed if no automatic landmarks are available (during initialization or when a user decides to delete all automatic landmarks). We observed that if the alignment fails, the matching also fails, and users rejected both. They then placed manual landmarks in combination with a few manual matches, which improved the next automatic matching computation without the automatic alignment. In combination with this workflow, we decided to always recompute the automatic landmarks after the matching for two main reasons. First, only matched MT ends should create landmarks, because the other MT ends possibly do not belong to each other. Second, during the manual matching, one can directly see the corresponding alignment changes due to the new landmark. This intuitive workflow combines local alignment and matching into a single control handle. Thus, the manual and automated matching can be applied iteratively to progress towards the best achievable stitching result. It was possible to create reasonable solutions even for cases where the fully automatic stitching process failed.

## Conclusions

In this paper, we have presented a tool to align serial tomograms in combination with the stitching of filamentous structures, specifically MTs in reconstructions of both mitotic and meiotic spindles. This tool allows users to visually validate the stitching process and support the automatic algorithms by placing manual landmarks and matches. Cases, in which the fully automatic workflow fails can now be solved with a reasonable investment of manual labor because a user can steer the whole process of stitching a stack of serial tomograms.

While stitching sections, minor errors of the tracing are often identified. For example, falsely traced MTs or erroneously connected MTs could be detected. Currently, it is not possible to edit the MT geometry with our tool. Users need to improve the MT tracing either before or after the stitching. It would be helpful to add such a functionality. This would allow users to immediately correct errors in the process of MT segmentation. For example, operations would be possible to add, split, remove, or merge MTs.

An open question is the computation or manual estimation of the gap between two adjacent sections. Currently, we expect that the user crops the tomograms in z-direction such that they fit seamlessly. It could be helpful to support setting the cropping regions interactively for each interface in the tool. This would probably improve the selection of cropping parameters, thus improving the overall alignment and matching procedure. It should be possible to compute good cropping parameters automatically from confirmed matches. In the future, this will need to be implemented in a version of the software package to come.

## Materials and methods

### Software availability and requirements

The described tool is available at https://www.zib.de/software/serial-section-aligner as an Amira custom module. To execute the tool, a commercial version of Amira (currently version 2019.2) is required. A free demo version can usually be obtained from Thermo Fisher Scientific upon request for a limited duration. As stated previously, performance tests were carried out on a desktop computer with an Intel Xeon i7 X5650 processor (6 cores) and an Nvidia GeForce GTX 1080 Ti graphics card. This hardware was sufficient to perform all interactive data processing at real time. 128 GB RAM were required to process the final stitching of the larger serial section tomogram stacks given in Table 1. If larger data sets are to be processed, it is recommended to increase the RAM. Moreover, a larger number of CPU cores could further improve processing performance.

### High-pressure freezing (HPF), freeze substitution (FS) and thin-layer embedding

Embryos, hermaphrodites and males and of wild-type *C. elegans* were prepared for high-pressure freezing as described previously [17–20]. HeLa cells were grown and frozen on Sapphire discs as reported [21]. Briefly, samples for the projects as listed in Table 1 were cryo-immobilized by using the following high-pressure freezers: EMPACT2+RTS (Leica Microsystems, Vienna, Austria), Leica EM ICE (Leica Microsystems, Vienna, Austria), HPF Compact 01 (M. Wohlwend, Sennwald, Switzerland). Freeze substitution was performed over 3 days at –90 °C in anhydrous acetone containing 1 % OsO_4_ and 0.1 % uranyl acetate using an automatic freeze substitution machine (EM AFS, Leica Microsystems, Vienna, Austria) Serial semi-thick sections (300nm) of Epon/Araldite-embedded samples were cut using an Ultracut UCT Microtome (Leica Microsystems, Vienna, Austria). Sections were collected on Formvar-coated copper slot grids and poststained with 2 % uranyl acetate in 70 % methanol followed by Reynold’s lead citrate.

### Electron tomography

Dual-axis electron tomography was performed as described [22]. Briefly, 20 nm-colloidal gold particles (Sigma-Aldrich) were attached to both sides of semi-thick sections collected on copper slot grids to serve as fiducial markers for subsequent image alignment. For electron tomography, series of tilted views were recorded using a TECNAI F30 transmission electron microscope (Thermo Fisher Scientific, Hillsboro, USA) operated at 300 kV. Images were captured every 1.0° over a ±60° range and a pixel size of 2.3 nm using a Gatan US1000 CCD camera (2k x 2k). For each serial section two montages were collected and combined to a supermontage using the IMOD software package [6] to cover the pole-to-pole distance of the spindles. For image processing the tilted views were aligned using the positions of the colloidal gold particles as fiducial markers. Tomograms were computed for each tilt axis using the R-weighted back-projection algorithm. For double-tilt data sets, montages of tomograms were aligned to each other and combined to a supermontage [22]. The number of tomograms and serial sections used for each presented data set are listed in Table 1.

## Acknowledgments

We are grateful to T. Fürstenhaupt (EM facility, Max Planck Institute of Molecular Cell Biology and Genetics, MPI-CBG) for continuous technical support. We also would like to thank I. Lantzsch, E. Szentgyörgyi and M. Kirchner for providing spindle data sets and comprehensive user feedback.

## Author Contributions

Conceptualization: NL, VJD, SP, TMR, DB.

Data curation: NL, GF, RK, SR.

Formal analysis: NL, FNB, VJD.

Funding acquisition: SP, TMR.

Investigation: NL, GF, RK, SR.

Methodology: NL, VJD, SP, DB.

Project administration: SP, TMR.

Resources: SP, TMR.

Software: NL, FNB, SP, DB.

Supervision: SP, DB.

Validation: GF, RK, SR.

Visualization: NL, SP, DB.

Writing – original draft: NL, SP.

Writing – review & editing: NL, VJD, FNB, SP, GF, RK, SR, TMR, DB.

## Video

**Video S1. Serial Section Aligner.** This video shows how the software tool described in this publication can be applied to stitch filamentous structures in image stacks from serial-section electron tomography. The video is subdivided into the following parts (times are given in parentheses): (1) Stack creation (00:07-00:29); (2) Navigation (00:30-01:46); (3) Alignment (01:47-02:46); (4) Matching (02:47-03:58); (5) Check matching (03:59-05:20); (6) Build stack (05:21-06:44).

